# Crinet: A computational tool to infer genome-wide competing endogenous RNA (ceRNA) interactions

**DOI:** 10.1101/2020.06.18.160408

**Authors:** Ziynet Nesibe Kesimoglu, Serdar Bozdag

## Abstract

To understand driving biological factors for complex diseases like cancer, regulatory circuity of genes needs to be discovered. Recently, a new gene regulation mechanism called competing endogenous RNA (ceRNA) interactions has been discovered. Certain genes targeted by common microRNAs (miRNAs) “compete” for these miRNAs, thereby regulate each other by making others free from miRNA regulation. Several computational tools have been published to infer ceRNA networks. In most existing tools, however, expression abundance sufficiency, collective regulation, and groupwise effect of ceRNAs are not considered. In this study, we developed a computational tool named Crinet to infer genome-wide ceRNA networks addressing critical drawbacks. Crinet considers all mRNAs, lncRNAs, and pseudogenes as potential ceRNAs and incorporates a network deconvolution method to exclude the spurious ceRNA pairs. We tested Crinet on breast cancer data in TCGA. Crinet inferred reproducible ceRNA interactions and groups, which were significantly enriched in the cancer-related genes and processes. We validated the selected miRNA-target interactions with the protein expression-based benchmarks and also evaluated the inferred ceRNA interactions predicting gene expression change in knockdown assays. The hub genes in the inferred ceRNA network included known suppressor/oncogene lncRNAs in breast cancer showing the importance of non-coding RNA’s inclusion for ceRNA inference. The source code of Crinet could be accessed on Github at https://github.com/bozdaglab/crinet.

## 1 Introduction

MicroRNAs (miRNAs) are small RNA types that regulate gene expression by binding to them. Recently, a new regulatory layer related to miRNAs has been discovered [25]: certain RNAs targeted by common miRNAs “compete” for these miRNAs and thereby regulate each other indirectly by making the other RNA(s) free from miRNA regulation. Such indirect interactions between RNAs are called competing endogenous RNA (ceRNA) interactions, which have important roles in diseases including cancer [16, 23, 30, 35]. There is a regulation multiplicity between miRNAs and RNAs, meaning that a miRNA could have multiple RNA targets, and an RNA could be targeted by multiple miRNAs. Given the enormous number of RNAs and difficulty of deciphering miRNA binding targets accurately, identifying ceRNA interactions experimentally is cost- and labor-prohibitive. Therefore, computational tools are crucial to infer ceRNA interactions in complex genomes like human. For the rest of the paper, “genes” refers to mRNAs, lncRNAs, and pseudogenes in our analysis.

Despite existing computational tools [5, 6, 10, 29], there exists crucial drawbacks. Current tools compute only pairwise ceRNA interactions or ceRNA modules, however, inferring groupwise ceRNA interactions should be considered since several ceRNAs could work together to sequester miRNA(s) targeting key ceRNA(s). Also, in the existing tools, a miRNA/gene could be assigned to many genes/miRNAs without considering the sufficiency of miRNA/gene expression abundance. Furthermore, since ceRNAs positively regulate each other, two genes having many common ceRNA partners might be inferred just because of the amplifying effect of regulation by common ceRNA partners. Thus, excluding these false positive ceRNA interactions is important.

In this study, we developed a computational tool named Crinet (CeRna Interaction NETwork) to infer genome-wide ceRNA interactions and groups to address the aforementioned drawbacks. To build our ceRNA network on a proper miRNA-target interaction set, we integrated expression datasets and binding scores for miRNA-target pairs considering expression abundance sufficiency. We computed ceRNA pairs considering strong regulation jointly. To cope with the spurious interactions, we excluded ceRNA pairs which were potentially inferred because they had significant overlapping of common ceRNA partners. We inferred ceRNA groups and integrated these groups into the ceRNA network. Getting the benefit of multiple biological datasets and including non-coding RNAs, this approach facilitates a better understanding of ceRNA regulatory mechanisms addressing important drawbacks, which would shed light on the underlying complex regulatory circuitry in disease conditions.

Crinet was applied to breast cancer dataset to infer ceRNA interactions and groups. We evaluated Crinet-selected miRNA-target interactions with protein expression-based benchmarks having increasing performance following filtering steps. Expression change in the inferred ceRNAs was highly affected following knockdown of their ceRNA partners. Inferred ceRNAs, hub ceRNAs, and some ceRNA groups were significantly associated with the cancer-related genes and processes, and consistently involved in the immune system related processes, thus Crinet-inferred ceRNAs could be important assets in the light of the studies confirming the relation between immunotherapy and cancer.

## 2 Materials and Methods

Crinet is a computational tool to infer genome-wide ceRNA interactions and groups (Fig. 1). Briefly, the first step is the data preparation step. In the second step, miRNA-target interactions are computed by incorporating expression datasets and considering expression abundance sufficiency. Starting with final miRNA-target interactions, ceRNA interactions are inferred in the third step. In the last step, ceRNA groups are inferred from the ceRNA network and integrated into the network. In the following, each step of Crinet is explained in more detail.

**Figure 1.**
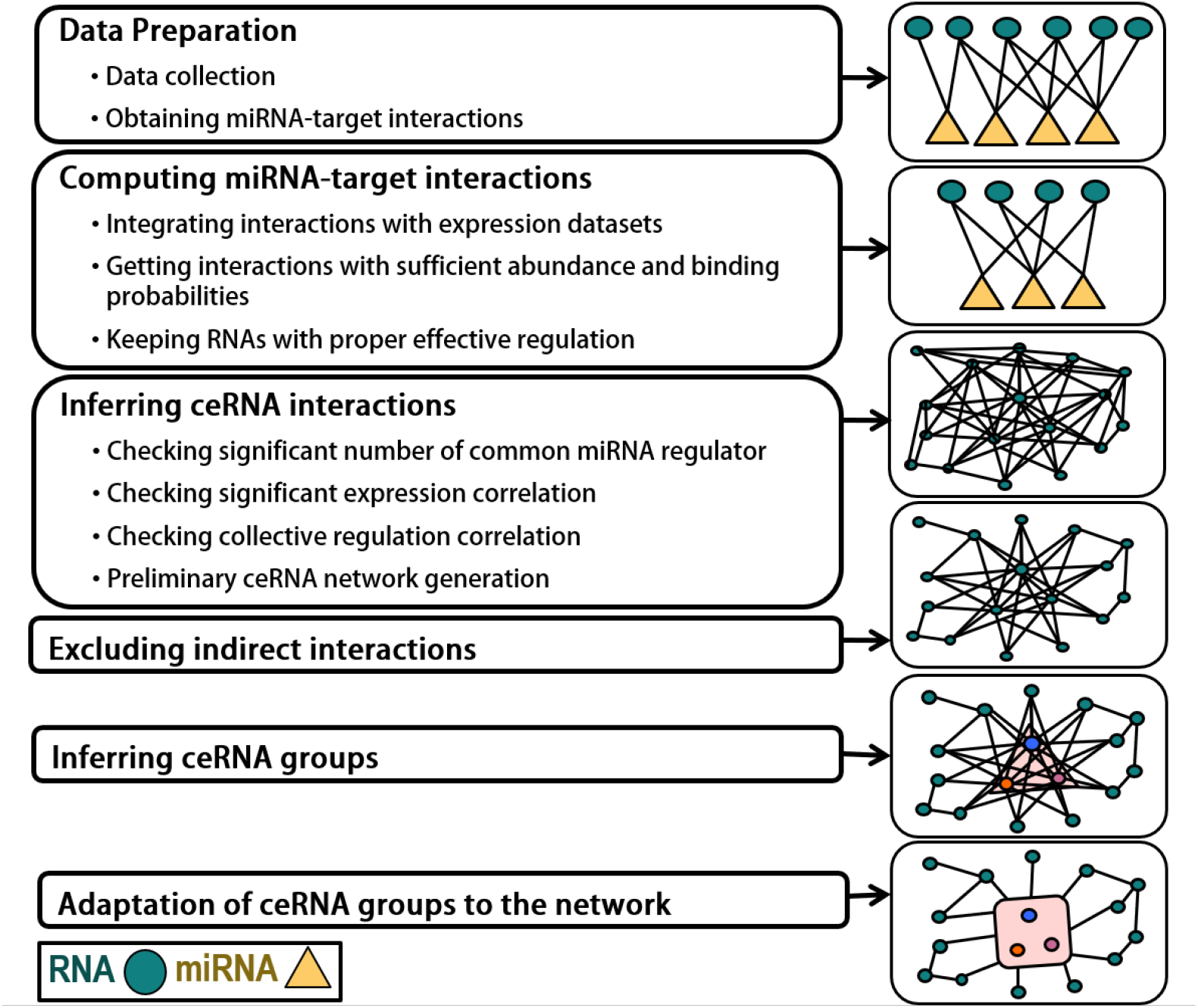
Crinet Pipeline

### 2.1 Data preparation

Crinet incorporates miRNA-target interactions with binding scores, gene-centric copy number aberration (CNA), and expression datasets. If binding scores are not available, the same score for all interactions could be used.

To collect datasets for Crinet, we used TCGAbiolinks R package [8] and obtained the datasets from the Cancer Genome Atlas (TCGA) project including gene expression, miRNA expression, and CNA for totally 1107 breast (BRCA) tumor samples (https://www.cancer.gov/tcga). We preprocessed each datatype separately obtaining normalized expression values (gene expression as FPKM and miRNA expression as RPM) and filtered lowly expressed genes (if FPKM *<* 1 and RPM = 0 for at least 15% samples). To get gene-centric CNA from a segmented dataset, we ran CNTools R package [33].

We obtained all conserved and nonconserved mRNA-target interactions with weighted context++ scores from TargetScan [1]. To compute pseudogene and lncRNA targets of miRNAs, we ran TargetScan separately for lncRNAs and pseudogenes, providing a constant dummy ORF length and supplying all transcript sequence as 3’UTR with the assumption that the entire sequence could be bound by miRNAs. If multiple scores exist for one pair due to the multiple transcripts, we used the strongest score.

Since there is a corresponding weighted context++ score in TargetScan results even for weak miRNA-target interactions, we kept the top 40% of ranked interactions with respect to weighted context++ scores. To fit different distributions of scores for mRNAs, lncRNAs, and pseudogenes, we combined z-normalized scores from each and applied min-max normalization after having all the scores in the range of -1 and 1. We used these normalized scores as the weight for each miRNA-target interaction assuming that these scores show the binding strength between miRNA and its gene target.

### 2.2 Computing miRNA-target interactions

In this step, we computed final miRNA-target interactions leveraging expression datasets and considering expression abundance sufficiency.

#### Integrating interactions with expression datasets (Correlation filtering)

Since miRNAs are known to repress their target genes, we kept only miRNA-target pairs having negative correlation between their expressions. By default, we used the correlation coefficient threshold of -0.1, which was the median correlation coefficients of all miRNA-target pairs. We also applied random sampling with replacement to compute correlation coefficient of each miRNA-target pair for 1000 times and required that the threshold was satisfied for ≥ 99% of the samplings.

#### Getting interactions with sufficient abundance and binding probabilities (Abundance filtering)

In the miRNA-target interaction sets, a miRNA could be assigned to mediate thousands of RNAs, and similarly an RNA could be assigned to many miRNAs as a potential target. To quantify the expression sufficiency for our putative interactions, we introduced *Interaction Regulation* (IR) formulated as:

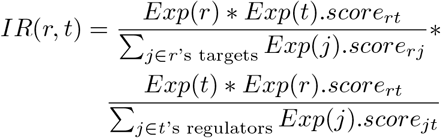

where *IR*(*r, t*) is the IR of the regulator *r* and the target *t* across samples, *Exp*(.) is the expression vector across samples, *score*_*rt*_ is the normalized binding score for the interaction between the regulator *r* and the target *t*. Using this formula, we kept the final miRNA-target interactions having high IRs (i.e., 80^th^ percentile of log of *IR*(*r, t*) *>* − 4.89, which was third quartile of all IRs through samples).

#### Keeping genes with proper effective regulation

To exclude genes from analysis if they were not under strong miRNA regulation based on the final miRNA-target interactions, we introduced *Effective Regulation* (ER) formulated as:

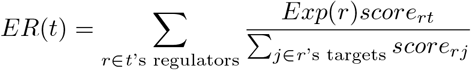

To keep genes with proper effective regulation by miRNAs, we filtered out genes without strong negative correlation (*<* −0.01) between its expression and ER, assuming that they did not have strong miRNA regulation for our specific dataset. We also applied random sampling with replacement to compute correlation for 1000 times and required that the threshold was satisfied for ≥99% of the samplings. We used the remained genes (called candidate genes) for further analysis.

### 2.3 Inferring ceRNA interactions

To infer ceRNA interactions, we generated all possible gene-gene combinations using candidate genes and filtered them based on the following criteria:

#### Checking significant number of common regulator

Since ceRNAs should have common miRNAs to compete for, we kept gene pairs having a significant number of common miRNA regulators (hypergeometric p-value *<* 0.01).

#### Checking significant expression correlation

Since the ceRNA pairs indirectly positively regulate each other and CNA considerably affects the expression values, we kept the gene pairs having significant partial correlation when excluding the CNA effect from the gene expression. By default, we used the correlation coefficient threshold of 0.55 (p-value *<* 0.01), which was the third quartile of the positive correlation (Supplementary Fig. 3).

#### Checking collective regulation

If there exists ceRNA regulation between two genes then both genes compete for common miRNAs and those miRNAs affect both genes simultaneously. We called this regulation *Collective Regulation* (CR) formulated as:

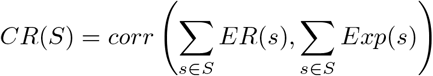

where *S* is a set of genes having ceRNA interactions and *corr*() is the Pearson correlation function. We kept the ceRNA pairs if they had a CR *<* − 0.01.

We applied random sampling with replacement for both partial correlation and CR measurements (last two steps) 100 times separately and kept the interactions when the threshold was satisfied for ≥ 99% of the samplings.

#### Excluding amplified interactions

Having common ceRNA partners between any two genes will increase the correlation between their expression. If a gene pair has too many common ceRNA partners, then some of the ceRNA interactions could be superior due to the high number of common ceRNA partners. To exclude such spurious interactions from our network, we employed a network deconvolution algorithm [11] and kept the top one-third of ranked interactions as our pairwise ceRNA network.

### 2.4 Inferring ceRNA groups

In ceRNA regulation, each ceRNA pair compete for common miRNA(s) and act as a decoy to make the other RNA free from miRNA regulation. However, this competition could occur among more than two RNAs, or between two groups of RNA. Based on this premise, Crinet inferred ceRNA groups in addition to ceRNA interactions.

To obtain ceRNA groups, we utilized one of the popular community detection algorithms named Walktrap [22] on the weighted ceRNA network where weights were normalized partial correlation coefficient. We kept the groups satisfying all the group conditions, otherwise split them iteratively. Three group requirements of Crinet are listed as follows:

1. **Common miRNA regulator:** To be able to compete for, all the group members were required to have at least one common miRNA regulator.
2. **Strong regulation effect:** CeRNAs in a group were expected to have a stronger miRNA regulation effect as a group than as individual ceRNAs. Thus, we required that CR of a group must be stronger (i.e. reduced) than the correlation between expression and ER of ≥ 90% of the group members. Moreover, to ensure that most of the group members would be under strong collective regulation effect, we required that an average difference between CR of the group and *corr*(*Exp*(*g*), *ER*(*g*)) for each gene *g* in the group was *>* 0.
3. **Compatibility with the network:**To hold inference consistency of the ceRNA network, for a given group, we called each ceRNA partner of group members as *neighbor* and expected the group to satisfy any two of the three conditions for at least 90% of its neighbors. The conditions are i) at least one common miRNA regulator between the group and the neighbor, ii) a strong Pearson correlation based on expression, and iii) strong collective regulation between the group and the neighbor.

## 3 Results

We tested Crinet on breast tumor samples from TCGA (Section 2.1 for details). We computed miRNA-target interactions (Table 1) and used them to infer 17,443 pairwise ceRNA interactions (Table 2). Using this pairwise ceRNA network, we obtained 81 ceRNA groups after applying 1508 iterations of Walktrap. Thirty five of these groups were connected to at least one node in the final network, while the others had interactions only within the group. After this step, we had our grouped ceRNA network with 4352 nodes (4317 individual genes and 35 groups of genes) and 17,274 edges between inferred nodes.

**Table 1.** Number of miRNA-target interactions after each miRNA-target interaction filtering step in Crinet. ALL: All obtained miRNA-target interactions; ALL40: Top 40% of ALL interactions based on weighted context++ score; CORR: After correlation-based filtering on ALL40 interactions (< −0:1); CORR+ABUN: After abundance-based filtering on CORR interactions (log IR > −4:89 for > 80% of samples)

| Step | # of Interactions | # of miRNAs | # of Genes |
| --- | --- | --- | --- |
| ALL | 8,193,904 | 888 | 28,129 |
| ALL40 | 3,285,523 | 888 | 27,999 |
| CORR | 535,284 | 888 | 23,099 |
| CORR+ABUN | 165,937 | 888 | 21,261 |

ALL: All obtained miRNA-target interactions; ALL40: Top 40% of ALL interactions based on weighted context++ score; CORR: After correlation-based filtering on ALL40 interactions ( $< -0.1$ ); CORR+ABUN: After abundance-based filtering on CORR interactions ( $\log IR > -4.89$ for $> 80\%$ of samples)

**Table 2.** Number of remained ceRNA pairs after each ceRNA interaction filtering step in Crinet. ALL: All candidate pairs after keeping proper genes with effective regulation; Step1: pairs with a significant overlap for common miRNAs; Step2: pairs after filtered based on partial correlation between gene expressions excluding copy number aberration effect; Step3: pairs after filtered based on collective regulation; Step4: pairs after applying random sampling for Step2 and Step3; Step5: pairs after applying the network deconvolution method to exclude the spurious ceRNA pairs.

| Step | # of Pairs | # of Genes |
| --- | --- | --- |
| ALL | 212,994,480 | 20,640 |
| Step1 | 16,082,000 | 20,640 |
| Step2 | 247,885 | 11,910 |
| Step3 | 209,220 | 11,726 |
| Step4 | 52,858 | 7,263 |
| Step5 | 17,443 | 4,494 |

To check the scale-free property and specificity of Crinet, we examined the inferred network (Supplementary Methods A.2 for details). Since biological networks generally exhibit scale-free property, we computed the inferred network’s degree probability distribution function. Our inferred ceRNA network had a negative slope with high fitness (*R*^2^ = 0.93), indicating that the inferred ceRNA network was scale-free (Supplementary Fig. 2). To evaluate the specificity of Crinet, we checked if our inferred ceRNA pairs existed in different regulatory layers, namely protein-protein interactions (PPIs) and transcription factor (TF)-gene interactions. We collected 1,663,810 TF-target interactions from TRRUST v2 [12] database and the ENCODE Transcription Factor Targets dataset [9], and 1,847,774 PPIs from BIOGRID v3.5.186 [26]. Within all inferred ceRNA interactions, very few interactions were TF-gene interactions (0.46%) and PPIs (0.51%) indicating that the regulatory relationships between inferred ceRNA interactions were not due TF or PPI effect.

**Figure 2.**
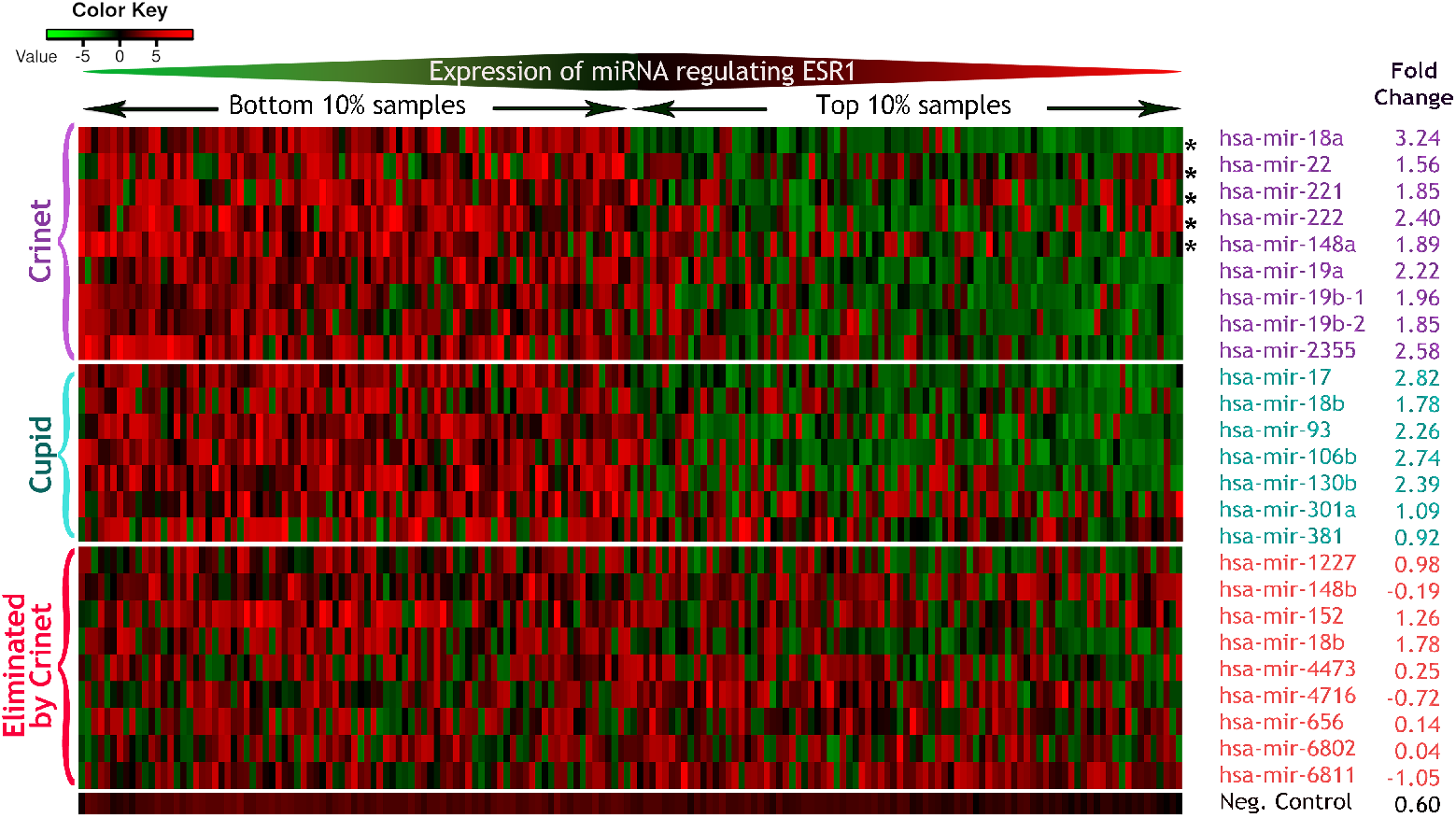
Heatmap showing protein expression of ESR1 for the top and bottom 10% ranked samples with respect to miRNA expression. Protein expression is shown for the top and bottom 10% samples ranked with respect to miRNA expression regulating ESR1 by Cupid-selected, Crinet-selected, Crinet-eliminated, and negative control along with the mean difference of log fold-change of protein expression for the bottom 10% with respect to the top 10% samples. Each row is independently ranked by miRNA expression. *Common miRNA regulators with Cupid

To check the reproducibility based on different datasets, the robustness to different hyperparameters, and the effect of each individual step in ceRNA inference, we conducted more detailed analysis of Crinet results (Supplementary Methods A.3 and A.4 for details). To check the reproducibility of Crinet based on different datasets, we ran Crinet on two equal-sized random samplings of the breast cancer dataset multiple times. To avoid bias in the comparisons, we ensured that both samplings had similar subtype distribution (namely Basal-like, Normal-like, Luminal-A, Luminal-B, and Her2-enriched). We observed highly overlapping interactions and ceRNAs among different runs (Supplementary Table 4). We checked the distribution of consistently overlapping ceRNAs and observed that the mean degree of these ceRNAs was much higher as compared to the overall mean degree (p-value *<* 2.10^−16^) suggesting that consistently inferred ceRNAs were the hub ceRNAs highly involving in our inferred ceRNA network. Moreover, to examine the effect of each step in Crinet, we disabled major steps in ceRNA inference and evaluated the results. Disabling individual steps made a substantial difference in the inferred results (Supplementary Table 1). However, when we modified the hyperparameters in each of these steps, we observed highly overlapping interactions (Supplementary Table 5 and Supplementary Table 6) suggesting that Crinet is robust to different choices of hyperparameters.

### 3.1 miRNA-target interaction filtering showed increasing performance on protein expression-based benchmarks

Since we built a ceRNA network relying on miRNA-target interactions, proper selection of these interactions is important; therefore, we evaluated each filtering step of miRNA-target interactions using protein expression-based benchmarks.

#### Transfection analysis

We utilized a Reverse Phase Protein Array (RPPA) dataset for MDA-MB-231 breast cancer cell line from The Cancer Proteome Atlas (TCPA) database (accession number: TCPA00000001) [2] to assess our miRNA-target interactions as in [5]. We used 104 antibodies, their fold-change for 141 transfected miRNAs, and mock controls. For each miRNA-target interaction, we measured expression fold ratio of each antibody of the target for the miRNA transfection relative to the average mock transfections. Table 3 confirms the preferential down-regulation of predicted miRNA targets, getting higher after each consecutive filtering step showing the positive effect of filtering for each independent interaction. We also checked the average of all targeting miRNAs per gene relative to average mock transfections and observed a similar down-regulation tendency. Although the ratio did not increase for the last step, it was due to few genes. The proteins ERCC1, BAK1, CTNNA1, PXN, MSH2, XIAP, MAPK3, EEF2K, CAV1, IGFBP2, PRKAA1, PCNA, CASP9, IGF1R, SMAD4, COL6A1, PIK3R1, CHEK1, EIF4E, PTK2, CDK1, SMAD1, BCL2L1, BCL2, LCK, DIABLO, NF2, and EIF4EBP1 some of which are known to be important in breast cancer were consistently down-regulated by their predicted miRNA regulators. As a negative control, we used non-inferred interactions and did not observe any strong down-regulation tendency for all the filtering steps for both phases (Supplementary Table 2).

**Table 3.** Evaluation of miRNA-target interaction filtering steps for the computed miRNA-target interactions using miRNA transfection data. Interaction phase shows the expression fold reduction of each antibody of target for its transfected miRNA regulator relative to mock transfection. Gene phase shows average expression fold reduction of each antibody of target for all transfected miRNAs regulators relative to mock transfection. Down-regulated over up-regulated numbers along with the ratio are shown (ratio is expected to be more than 1 to have down-regulation tendency. Higher is better). See Table 1’s caption for the definition of row labels.

| Step | # of Interactions | Interaction Phase | Gene Phase |
| --- | --- | --- | --- |
| ALL | 8,193,904 | 6,248/4,932=1.3 | 90/74=1.2 |
| ALL40 | 3,285,523 | 613/403=1.5 | 97/59=1.6 |
| CORR | 535,284 | 183/112=1.6 | 71/37=1.9 |
| CORR+ABUN | 165,937 | 179/111=1.6 | 68/39=1.7 |

#### Protein expression anticorrelation analysis

Using protein expression dataset from TCPA matching with breast tumor samples in our analysis, we analyzed the negative correlation between miRNA expression and protein expression of their targets for each applicable miRNA-target interaction.

While the anticorrelation tendency slightly decreases after selecting top 40% miRNA-target interactions, our filtering steps substantially increased the ratios showing the anticorrelation tendency in our selected interactions (Table 4). Also, while considering average miRNAs per target, we had a slight anticorrelation tendency for top 40% interactions; however, our filtering steps increased the anticorrelation tendency much more. As negative control, non-inferred interactions did not show strong tendency, and even the tendency was towards to the up-regulation for the gene phase (Supplementary Table 3).

**Table 4.** Evaluation of miRNA-target interaction filtering steps for the computed miRNA-target interactions based on miRNA-protein expression anticorrelation. Interaction phase shows the anticorrelated expressions of miRNA-protein target pairs with respect to positively correlated pairs. Gene phase shows the anticorrelated expression of target with average of all miRNA regulators with respect to the one having positive correlation. (ratio is expected to be more than 1 to have anticorrelation tendency. Higher is better). See Table 1’s caption for the definition of row labels.

| Step | # of Interactions | Interaction Phase | Gene Phase |
| --- | --- | --- | --- |
| ALL | 8,193,904 | 38,505/33,243=1.2 | 95/106=0.9 |
| ALL40 | 3,285,523 | 2,720/2,688=1.0 | 107/91=1.2 |
| CORR | 535,284 | 751/370=2.0 | 115/50=2.3 |
| CORR+ABUN | 165,937 | 719/352=2.0 | 114/47=2.4 |

#### Protein expression analysis with ESR1

To evaluate predicted miRNA-target interactions in [5], the authors focused on the ESR1 protein, showing that ESR1 protein expression in TCGA breast cancer tumors (profiled by RPPA using the antibody ER.alpha.R.V GBL.9014870) had a strong negative correlation with the expression of predicted miRNA regulators. Ranking samples based on miRNA expression, the top 10% and bottom 10% samples were compared based on ESR1 protein expression. Similarly, we generated a heatmap showing protein expression for Crinet-selected miRNAs regulating ESR1. Fig. 2 shows nine Crinet-selected and 12 Cupid-selected miRNAs regulating ESR1, having five miRNAs as common. We quantified the anticorrelation between miRNA and protein expression by measuring fold-change of mean protein expression for the top 10% samples with respect to the bottom 10%. Our results indicated that the expression of Crinet-selected miRNAs for ESR1 had high anticorrelation with protein expression with high fold-change consistently while Cupid had some low fold-change such as hsa-mir-381.

Fig. 2 also illustrates nine regulators eliminated by Crinet following expression correlation and sufficient abundance filtering of miRNA-target selection. These interactions did not show strong anticorrelation between miRNA and protein expression with respect to fold-change for the majority of miRNAs, showing the strength of Crinet filtering approach. As a negative control, we added the average of 100 random miRNAs which were not selected as ESR1’s regulator by Cupid and Crinet, and they exhibited a low fold-change.

### 3.2 Inferred ceRNA interactions were able to predict gene expression change

To assess the accuracy for ceRNA inference, we used the Library of Integrated Network-based Cellular Signature (LINCS) [18] L1000 shRNA-mediated gene knockdown experiment in breast cancer cell line as in [6] and checked whether ceRNA interactions can predict the effects of RNAi-mediated gene silencing perturbations in MCF7 cells. Since Crinet starts with a high number of genes, it was not computationally feasible to run many tools with our dataset. However, Hermes [6] runs any given ceRNA pair independently, therefore we ran Hermes for the genes in the knockdown assays using the same expression datasets and Crinet-selected miRNA-target interactions. LINCS database is a rich resource having an expression change of nearly 1000 genes as a response to a silenced gene. When a gene is silenced then its ceRNA partner will be affected since more miRNA regulators will be available to suppress the ceRNA partner. Thus, given a ceRNA pair, expression level should be lower in response to the silenced ceRNA partners in comparison to the genes that are not ceRNA partners. Based on this assumption, we evaluated the Crinet- and Hermes-inferred networks. The accuracy of this assessment is shown in Table 5. Since Hermes was not selective in terms of the number of ceRNA interactions by inferring many significant interactions, we evaluated several networks from Hermes till having similar number of genes with Crinet in the knockdown assessment. Based on these results, Crinet outperformed Hermes at predicting gene expression change of ceRNA partners for each timepoint and for overall accuracy.

**Table 5.** Evaluation of the accuracy of Crinet- and Hermes-inferred ceRNA interactions based on the shRNA-mediated gene knockdown experiment. Analysis to check the accuracy of inferred ceRNA interactions using LINCS-L1000 shRNA-mediated gene knockdown experiment in breast cancer cell line. Based on the ratios of gene expression fold-change following the knockdown of its ceRNA partners to following the genes that are not its ceRNA partners for each perturbagen ceRNA, the accuracy of a ceRNA network was accepted as the percentage of ceRNAs whose ratios were smaller than 1 with respect to all ceRNAs. We calculated the accuracy separately for each different timepoint (96h & 144h) and combined timepoints as the overall. Hermes’s x.run had 10^(*x*+1)^ for the significance of the common miRNA size and 10^*x*+2)^ for the significance of conditional regulation.

|  | 96h Timepoint | 144h Timepoint | Overall |
| --- | --- | --- | --- |
| Crinet | 48/77 = 62% | 44/77 = 57% | 92/154 = 60% |
| Hermes 1.run | 412/897 = 46% | 441/913 = 48% | 853/1810 = 47% |
| Hermes 2.run | 199/466 = 43% | 217/481 = 45% | 416/947 = 44% |
| Hermes 3.run | 109/269 = 41% | 121/273 = 44% | 230/542 = 42% |
| Hermes 4.run | 65/143 = 45% | 68/144 = 47% | 133/287 = 46% |
| Hermes 5.run | 33/69 = 48% | 39/71 = 55% | 72/140 = 51% |

### 3.3 Inferred ceRNAs were significantly associated with the known cancer genes and cancer-related processes

To analyze the biological significance of the inferred ceRNA network, we applied enrichment analysis for the inferred ceRNAs. We used ClusterProfiler R package [32] for all enrichment analysis. The inferred ceRNAs were significantly enriched in 398 GO terms from biological process ontology and 39 KEGG pathways. To associate enriched terms to broader categories, we analyzed GO Slim terms (Supplementary Table 8). Inferred ceRNAs were mostly involved in biological processes including immune system process, cell differentiation, cell death, cell cycle, response to stress, and cell-cell signaling. These suggest that ceRNA interactions could have important role in biological processes in cancer.

To check if the inferred ceRNAs were associated with the cancer-related genes, we collected 3078 known cancer genes obtained from Cancer Gene Census in COSMIC v91 [27], Bushman’s cancer gene list v3 [17], human oncogenes from ONGene [20], Network of Cancer Genes 6.0 [24], and LncRNADisease database from the Cui Lab [4]. In our inferred ceRNAs, we had significant overlap (hypergeometric test p-value 9.10^−105^) between the known cancer genes and the inferred ceRNAs having 789 out of 3078 known cancer genes in our inferred network. When we repeated the same analysis for non-inferred genes, they did not show significant p-value (almost 1). We also had 54 breast cancer-related genes from LncRNADisease and Network of Cancer Genes 6.0 databases, 14 out of 54 breast cancer-related genes were inferred in our network. These results indicated that inferred ceRNAs were significantly associated with known cancer genes.

Moreover, we analyzed the hub ceRNAs in our network. The degree distribution of our network had a median of three with a maximum of 152 and a minimum of 1. We got the top 81 ceRNAs having a degree of 50 or more in our network as hub genes. The hub genes were significantly enriched in 58 GO terms and eight KEGG pathways. We investigated the GO Slim terms from biological processes ontology for hub genes, and enriched terms included immune system process, cell death, cell differentiation, cell motility, and cell cycle. Also, these hub genes were among the known cancer genes with a hypergeometric p-value of 0.0009. Specifically, 15 out of 81 genes were involved in the known cancer genes, while three of them were breast cancer-related. These suggest that hub ceRNAs in the inferred network were involved in the important biological processes in cancer.

Among hub genes were some lncRNAs with known involvement in cancer. For instance, MAGI2-AS3 had a degree of 89 being highly connected for ceRNA regulation in our network, and it is known as suppressor involving in cell growth [31]. MALAT1, which had a degree of 51, contributes significantly to cancer initiation and progression in breast cancer [15]. Some other lncRNAs had also important functionality: MIR100HG as an oncogene involving in proliferation [28], ITGB2-AS1 as an oncogene involving migration and invasion [19], and MEG3 as a suppressor involving in proliferation and EMT [34].

### 3.4 Inferred ceRNA groups included known cancer-related genes and were enriched in cancer-related processes

To evaluate the inferred ceRNA groups, we performed enrichment for four of 81 inferred ceRNA groups that have more than three members (Supplementary Fig. 4). CeRNA groups had a significant overlap with the cancer-related genes (hypergeometric p-value *<* 0.0006). Group 1 genes were enriched with 82 GO terms from biological process ontology and 5 KEGG pathways, while group 3 were enriched with 35 GO terms and group 4 with 17 GO terms. Moreover, group 1 was highly enriched with GO Slim terms including cell cycle, response to stress, DNA metabolic process, chromosome segregation, and cell division. Although group 2 had limited mRNAs, all the remaining groups (groups 2, 3, and 4) were consistently enriched with GO Slim term immune system process. Additionally, group 3 had GO Slim terms including cell death, cell adhesion, and cell motility, while group 4 included response to stress (Supplementary Table 7 for details). CeRNA groups were significantly overlapped with known cancer-related genes and significantly enriched in biological processes suggesting that ceRNA groups could have important roles as a group in cancer including the immune system and cell repair.

## 4 Discussion

In this study, we developed a computational tool named Crinet to infer pairwise and groupwise ceRNA interactions and applied it to the breast tumor samples. Leveraging multiple types of biological datasets, considering expression abundance between miRNA and their targets, and excluding amplifying effect of ceRNA regulation, we inferred a ceRNA network including 17,274 ceRNA interactions between 4352 ceRNAs/ceRNA groups.

Unlike the existing tools, we filtered the miRNA-target interactions considering abundance sufficiency and binding scores. We introduced Interaction Regulation (IR) score, and confirmed that the miRNAs with the highest number of targets and low expression levels were successfully filtered out (Supplementary Methods A.1 and Supplementary Fig. 1).

Since ceRNAs positively regulate each other, expecting positive correlation between expression of ceRNAs is a common approach [25]. While some studies use additional metrics in addition to simply calculating correlation [10, 21], we measured the partial correlation excluding the CNA effect since expression values are highly affected by copy number amplification and deletion. Different from the studies that analyze only differentially expressed (DE) genes to have a computationally manageable number, we included all lncRNAs and pseudogenes in addition to all coding RNAs (including non-DE mRNAs) to understand comprehensive ceRNA regulation. We had the known lncRNA oncogenes and suppressors as highly connected in the inferred ceRNA network indicating that the lncRNAs could have an important role in ceRNA regulation. When the expression profiling of other data types such as circular RNAs is available, including them into the pipeline could be worthy of further investigation to improve ceRNA inference [14]. We also started with a large set of miRNA-target interactions since we utilized weighted context++ score as a proxy for binding probability. This enabled us to include all possible gene-gene interactions based on common regulators especially for the measurements like our collective and effective regulation. Our carefully designed miRNA-target interaction filtering steps were able to eliminate high likely false-positive interactions confirmed by the protein expression-based benchmarks.

We used a network deconvolution method [11] and eliminated the amplifying effect of ceRNA pairs, which was not addressed by the previous studies. Crinet inferred ceRNA groups holding the inference consistency in the ceRNA network. In that way, we did not find only a group of closely related genes, but we had a ceRNA network in which the groups and individual ceRNAs had ceRNA interactions showing the comprehensive regulation.

Delving into the biological significance, inferred ceRNAs, ceRNA groups, and hub ceRNAs were significantly enriched in the known cancer-related genes and processes suggesting that ceRNAs could serve important processes in cancer. We consistently had immune system process as the significantly enriched GO term for the inferred ceRNAs, some ceRNA groups, and the hub ceRNAs. There are studies confirming that the weakness in the immune system function is closely related to tumorigenesis [3]. In [13], authors investigated ceRNA networks in Papillary Thyroid Carcinoma and disclosed the combined regulation of immune responses from these networks. In [7], authors unraveled the prognostic significance of ceRNA interactions among immune response genes in glioblastoma multiforme. Considering the immunotherapy as an emerging field in cancer with its potential to provide a strong response in cancer patients, novel Crinet-inferred ceRNA interactions and groups significantly enriched in immune-related processes could be important assets.

## Acknowledgments

This work was supported by the National Institute of General Medical Sciences of the National Institutes of Health under Award Number R35GM133657. Supplementary material referred in this manuscript can be accessed at the preprint version on bioRxiv at https://doi.org/10.1101/2020.06.18.160408.

## Notes

https://github.com/bozdaglab/crinet

### Competing Interest Statement

The authors have declared no competing interest.

## References

1. V. Agarwal, G. W. Bell, J.-W. Nam, and D. P. Bartel. Predicting effective microrna target sites in mammalian mrnas. elife, 4:e05005, 2015.

2. R. Akbani, P. K. S. Ng, H. M. Werner, M. Shahmoradgoli, F. Zhang, Z. Ju, W. Liu, J.-Y. Yang, K. Yoshihara, J. Li, et al. A pan-cancer proteomic perspective on the cancer genome atlas. Nature communications, 5(1):1–15, 2014.

3. A. B. Bigley, G. Spielmann, E. C. LaVoy, and R. J. Simpson. Can exercise-related improvements in immunity influence cancer prevention and prognosis in the elderly? Maturitas, 76(1):51–56, 2013.

4. G. Chen, Z. Wang, D. Wang, C. Qiu, M. Liu, X. Chen, Q. Zhang, G. Yan, and Q. Cui. Lncrnadisease: a database for long-non-coding rna-associated diseases. Nucleic acids research, 41(D1):D983–D986, 2012.

5. H.-S. Chiu, D. Llobet-Navas, X. Yang, W.-J. Chung, A. Ambesi-Impiombato, Iyer, H. R. Kim, E. G. Seviour, Z. Luo, V. Sehgal, et al. Cupid: simultaneous reconstruction of microrna-target and cerna networks. Genome research, 25(2):257–267, 2015.

6. H.-S. Chiu, M. R. Martínez, M. Bansal, A. Subramanian, T. R. Golub, X. Yang, P. Sumazin, and A. Califano. High-throughput validation of cerna regulatory networks. BMC genomics, 18(1):418, 2017.

7. Y.-C. Chiu, L.-J. Wang, T.-P. Lu, T.-H. Hsiao, E. Y. Chuang, and Y. Chen. Differential correlation analysis of glioblastoma reveals immune cerna interactions predictive of patient survival. BMC bioinformatics, 18(1):132, 2017.

8. A. Colaprico, T. C. Silva, C. Olsen, L. Garofano, C. Cava, D. Garolini, T. S. Sabedot, T. M. Malta, S. M. Pagnotta, I. Castiglioni, et al. Tcgabiolinks: an r/bioconductor package for integrative analysis of tcga data. Nucleic acids research, 44(8):e71–e71, 2016.

9. E. P. Consortium et al. A user’s guide to the encyclopedia of dna elements (encode). PLoS biology, 9(4), 2011.

10. D. Do and S. Bozdag. Cancerin: A computational pipeline to infer cancer-associated cerna interaction networks. PLoS computational biology, 14(7):e1006318, 2018.

11. S. Feizi, D. Marbach, M. Médard, and M. Kellis. Network deconvolution as a general method to distinguish direct dependencies in networks. Nature biotechnology, 31(8):726, 2013.

12. H. Han, J.-W. Cho, S. Lee, A. Yun, H. Kim, D. Bae, S. Yang, C. Y. Kim, M. Lee, E. Kim, et al. Trrust v2: an expanded reference database of human and mouse transcriptional regulatory interactions. Nucleic acids research, 46(D1):D380–D386, 2018.

13. C.-T. Huang, Y.-J. Oyang, H.-C. Huang, and H.-F. Juan. Microrna-mediated networks underlie immune response regulation in papillary thyroid carcinoma. Scientific reports, 4:6495, 2014.

14. M. Huang, Z. Zhong, M. Lv, J. Shu, Q. Tian, and J. Chen. Comprehensive analysis of differentially expressed profiles of lncrnas and circrnas with associated co-expression and cerna networks in bladder carcinoma. Oncotarget, 7(30):47186, 2016.

15. J. Kim, H.-L. Piao, B.-J. Kim, F. Yao, Z. Han, Y. Wang, Z. Xiao, A. N. Siverly, S. E. Lawhon, B. N. Ton, et al. Long noncoding rna malat1 suppresses breast cancer metastasis. Nature genetics, 50(12):1705–1715, 2018.

16. M. S. Kumar, E. Armenteros-Monterroso, P. East, P. Chakravorty, N. Matthews, M. M. Winslow, and J. Downward. Hmga2 functions as a competing endogenous rna to promote lung cancer progression. Nature, 505(7482):212–217, 2014.

17. B. Lab. Bushman lab: Cancer gene list (version 4). Available from: http://www.bushmanlab.org/links/genelists, 05 2018. Accessed: 2020-06-06.

18. C. Liu, J. Su, F. Yang, K. Wei, J. Ma, and X. Zhou. Compound signature detection on lincs l1000 big data. Molecular BioSystems, 11(3):714–722, 2015.

19. M. Liu, L. Gou, J. Xia, Q. Wan, Y. Jiang, S. Sun, M. Tang, T. He, and Y. Zhang. Lncrna itgb2-as1 could promote the migration and invasion of breast cancer cells through up-regulating itgb2. International journal of molecular sciences, 19(7):1866, 2018.

20. Y. Liu, J. Sun, and M. Zhao. Ongene: A literature-based database for human oncogenes. J Genet Genomics, 44(2):119–121, 2017.

21. P. Paci, T. Colombo, and L. Farina. Computational analysis identifies a sponge interaction network between long non-coding rnas and messenger rnas in human breast cancer. BMC systems biology, 8(1):83, 2014.

22. P. Pons and M. Latapy. Computing communities in large networks using random walks. In International symposium on computer and information sciences, pages 284–293. Springer, 2005.

23. X. Qi, D.-H. Zhang, N. Wu, J.-H. Xiao, X. Wang, and W. Ma. cerna in cancer: possible functions and clinical implications. Journal of Medical Genetics, 52(10):710–718, 2015.

24. D. Repana, J. Nulsen, L. Dressler, M. Bortolomeazzi, S. Kuppili Venkata, Tourna, A. Yakovleva, T. Palmieri, and F. Ciccarelli. The network of cancer genes (ncg): a comprehensive catalogue of known and candidate cancer genes from cancer sequencing screens. Genome Biology, 20, 01 2019.

25. L. Salmena, L. Poliseno, Y. Tay, L. Kats, and P. P. Pandolfi. A cerna hypothesis: the rosetta stone of a hidden rna language? Cell, 146(3):353–358, 2011.

26. C. Stark, B.-J. Breitkreutz, T. Reguly, L. Boucher, A. Breitkreutz, and M. Tyers. Biogrid: a general repository for interaction datasets. Nucleic acids research, 34(suppl 1):D535–D539, 2006.

27. J. G. Tate, S. Bamford, H. C. Jubb, Z. Sondka, D. M. Beare, N. Bindal, H. Boutselakis, C. G. Cole, C. Creatore, E. Dawson, P. Fish, B. Harsha, C. Hathaway, S. C. Jupe, C. Y. Kok, K. Noble, L. Ponting, C. C. Ramshaw, C. E. Rye, H. E. Speedy, R. Stefancsik, S. L. Thompson, S. Wang, S. Ward, P. J. Campbell, and S. A. Forbes. COSMIC: the Catalogue Of Somatic Mutations In Cancer. Nucleic Acids Research, 47(D1):D941–D947, 10 2018.

28. S. Wang, H. Ke, H. Zhang, Y. Ma, L. Ao, L. Zou, Q. Yang, H. Zhu, J. Nie, C. Wu, et al. Lncrna mir100hg promotes cell proliferation in triple-negative breast cancer through triplex formation with p27 loci. Cell death & disease, 9(8):1–11, 2018.

29. X. Wen, L. Gao, and Y. Hu. Lacemodule: Identification of competing endogenous rna modules by integrating dynamic correlation. Frontiers in genetics, 11:235, 2020.

30. J. Yang, T. Li, C. Gao, X. Lv, K. Liu, H. Song, Y. Xing, and T. Xi. Foxo1 3 utr functions as a cerna in repressing the metastases of breast cancer cells via regulating mirna activity. FEBS letters, 588(17):3218–3224, 2014.

31. Y. Yang, H. Yang, M. Xu, H. Zhang, M. Sun, P. Mu, T. Dong, S. Du, and K. Liu. Long non-coding rna (lncrna) magi2-as3 inhibits breast cancer cell growth by targeting the fas/fasl signalling pathway. Human cell, 31(3):232–241, 2018.

32. G. Yu, L.-G. Wang, Y. Han, and Q.-Y. He. clusterprofiler: an r package for comparing biological themes among gene clusters. Omics: a journal of integrative biology, 16(5):284–287, 2012.

33. J. Zhang. Cntools: Convert segment data into a region by sample matrix to allow for other high level computational analyses. R package (Version 1.6. 0.), 2016.

34. W. Zhang, S. Shi, J. Jiang, X. Li, H. Lu, and F. Ren. Lncrna meg3 inhibits cell epithelial-mesenchymal transition by sponging mir-421 targeting e-cadherin in breast cancer. Biomedicine & Pharmacotherapy, 91:312–319, 2017.

35. M. Zhou, X. Wang, H. Shi, L. Cheng, Z. Wang, H. Zhao, L. Yang, and J. Sun. Characterization of long non-coding rna-associated cerna network to reveal potential prognostic lncrna biomarkers in human ovarian cancer. Oncotarget, 7(11):12598, 2016.

