## Supplemental Material for "Crinet: A computational tool to infer genome-wide competing endogenous RNA (ceRNA) interactions"

**1 Marquette University, Department of Computer Science, Milwaukee, WI, USA**

### Abstract

To understand driving biological factors for complex diseases like cancer, regulatory circuity of genes needs to be discovered. Recently, a new gene regulation mechanism called competing endogenous RNA (ceRNA) interactions has been discovered. Certain genes targeted by common microRNAs (miRNAs) "compete" for these miRNAs, thereby regulate each other by making others free from miRNA regulation. Several computational tools have been published to infer ceRNA networks. In most existing tools, however, expression abundance sufficiency, collective regulation, and groupwise effect of ceRNAs are not considered. In this study, we developed a computational tool named Crinet to infer genome-wide ceRNA networks addressing critical drawbacks. Crinet considers all mRNAs, lncRNAs, and pseudogenes as potential ceRNAs and incorporates a network deconvolution method to exclude the spurious ceRNA pairs. We tested Crinet on breast cancer data in TCGA. Crinet inferred reproducible ceRNA interactions and groups, which were significantly enriched in the cancer-related genes and processes. We validated the selected miRNA-target interactions with the protein expression-based benchmarks and also evaluated the inferred ceRNA interactions predicting gene expression change in knockdown assays. The hub genes in the inferred ceRNA network included known suppressor/oncogene lncRNAs in breast cancer showing the importance of non-coding RNA's inclusion for ceRNA inference. The source code of Crinet could be accessed on Github at https://github.com/bozdaglab/crinet.

### A Supplementary Methods

#### A.1 Our novel scoring for sufficiency of interactions drastically dropped the target distribution of miRNAs.

In this section, we evaluated our novel Interaction Regulation (IR) score that we used to eliminate miRNA-target interactions if there is no sufficient expression for miRNA or target. Since we had a much smaller number of miRNA as compared to targets, this step will affect the miRNAs more in terms of a decreased number of interactions. If a miRNA has many targets but low abundance, we expected a drastic change in the number of targets of this miRNA. To analyze the number of decrease in target numbers, we got the top 25% miRNAs (222 out of 888 miRNAs) with the highest number of targets in miRNA-target interactions set before abundance filtering and after abundance filtering. As in Figure 1A, it is clear that log median expression of top miRNAs are much higher after abundance filtering, suggesting that our scoring works well to eliminate the interactions with many targets without sufficient abundance.

Overall, the median of top 25% miRNAs with a high number of targets before abundance filtering had a minimum of 877 targets and maximum of 2428 targets with median 1155.5, however, the log median expression of these miRNAs had a median of 0.42 (having minimum 0 and maximum 67484). On the other side, after abundance filtering, top miRNAs with the highest number of targets had a minimum 222 and a maximum of 1333 targets with 431 as the median. With abundance filtering, naturally, the overall target numbers are dropped with the same number of miRNAs kept. However, the log median expression for top 25% miRNAs had a median of 45.74 (having minimum 0 and maximum 251417), much higher as compared to before abundance filtering. As expected from our novel scoring, miRNAs had a high number of targets if their abundance was sufficient.

Specifically, we got the top 6 miRNAs with the highest number of targets before the abundance step shown in Figure 1A. Namely, hsa-mir-939 with 2428 targets, hsa-mir-628 with 1994 targets, hsa-mir-4726 with 1978 targets, hsa-mir-616 with 1971 targets, hsa-mir-3191 with 1884 targets, and hsa-mir-4677 with 1806 targets. However, the median of these miRNAs are pretty low (0.9, 31.8, 0.0, 3.1, 0.2, 8.7, respectively). When we checked their target numbers after abundance filtering, they significantly dropped to 519, 729, 300, 424, 218, and 692 respectively. Similarly, when we got the top 6 miRNAs with the highest targets after abundance filtering, we had hsa-mir-22 with 1333 targets, hsa-mir-142 with 1323 targets, hsa-mir-210 with 1281 targets, hsa-mir-148b with 1233 targets, hsa-mir-17 with 1176 targets, and hsa-mir-183 with 1173 targets. When we checked their median miRNA expression, they have 67484.1, 1899.9, 385.1, 186.8, 465.7, and 15124.2 respectively. These expression values are high enough, especially when compared to top ones before abundance filtering. When we checked the target numbers before abundance filtering, they are still high with the numbers 1641, 1551, 1694, 1783, 1376, and 1445, respectively. These analyses showed that our scoring works well to exclude miRNA-target interactions from interactions set considering the sufficiency of abundance.

Moreover, we analyzed the number of targets overall before and after abundance filtering. Before the abundance step, we had miRNAs having targets starting from 8 to 2428 with a median of 519. Even keeping all the miRNAs in the set after abundance filtering, the target number range was starting from 6 to 1333 with a median 99.5 (Figure 1B).

Also, we analyzed the number of reduced targets for each miRNA after abundance filtering. When we just considered the miRNAs that are reducing more than half of their targets after abundance filtering, their number of targets had a median of 591. When we checked the median target number for the miRNAs reducing **more than** 75% of their targets, it increased to 622 targets per miRNA (with minimum 39 and maximum 2,428), while it dropped to 305 (with minimum 8 and maximum 1,694) for miRNAs reducing **less than** 25% of their targets. This shows the miRNAs with a higher number of targets reducing their targets even as a percentage, showing the preference is not because that they have a higher number of targets, but the ratio of reduced target number relative to all targets before abundance filtering is also high for them.

We also observed that the density plot showed the density of the reduced number of miRNA targets with a peak occurring around 0, is much lower than the expected reduction for any miRNA, which is 415.9 (mean of overall reduced target number per miRNA with minimum 0 and maximum 1,909) (Figure 1C). These analyses suggested that we eliminated miRNA-target interactions having low abundance and many targets applying our novel scoring to the miRNA-target interaction set.

**Density of ceRNA network was slightly more sparse after losing many genes following network deconvolution filtering:** Also, we investigated the density of the final network as compared to before network deconvolution (ND) step.

Since we applied ND to eliminate the amplifying effect of ceRNA interactions, we expected to have a more sparse network, even we expected to lose some ceRNAs totally from the network. We examined the remained network's edge density (the ratio of the number of edges and the number of possible edges) and transitivity (the probability that the adjacent nodes of a node are connected, also called the clustering coefficient). When we applied ND filtering, degree distribution in the final network was significantly lower as compared to the network before applying ND (p-value $< 10^{-15}$). Even we lost almost 3,000 ceRNAs from the network after applying the ND method, we had lower degree distribution, lower transitivity (dropped from 0.22 to 0.14), and slightly lower edge density (dropped from 0.0020 to 0.0017) for our final network.

### A.2 Inferred network was scale-free as a biological network and specific to Crinet

Since biological networks generally exhibit scale-free property, we checked whether our inferred ceRNA network is scale-free by computing its degree probability distribution function. Following the power-law rule, we fitted linear regression for the log of ceRNA's degree probability to the log of ceRNA's degree. The resulting plot of the inferred ceRNA network (Figure 2) had a negative slope with high fitness ($R^2 = 0.93$), indicating that the inferred ceRNA network was scale-free.

To show the specificity of Crinet, we compared our inferred ceRNA pairs from different regulatory layers, namely protein-protein interactions (PPIs) and transcription factor (TF)-gene interactions. We collected 1,663,810 TF-target interactions from TRRUST v2 [3] database and from the ENCODE Transcription Factor Targets dataset [1]. Within all inferred ceRNA interactions, very few interactions were TF-gene interactions (0.46% of all inferred interactions). Similarly, we collected 1,847,774 PPIs from BIOGRID v3.5.186 [4]. We also found that very few interactions were also PPIs (0.51% of all inferred interactions).

### A.3 Results were reproducible and robust to hyperparameter selection

To check the reproducibility of Crinet based on different datasets and hyperparameters, we ran whole pipeline multiple times with different datasets and compared the overlapping of the inferred networks. In addition to that, we examined that our pipeline is robust to hyperparameter selection running the whole pipeline with different thresholds for each step, and checking the overlapping ceRNA pairs.
**Reproducibility of Crinet:** We ran Crinet separately for randomly selected two different breast tumor sample sets. We divided all the breast tumor samples we had in our dataset into two equal-sized random sets with similar subtype distribution (namely basal-like, normal-like, luminal-A, luminal-B, Her2-enriched) to avoid bias in results. We ran the pipeline for both subsets and obtained inferred ceRNA interactions. We repeated this process for other random subsets of our samples, again having approximately the same number of samples for subtypes. We compared the results of two runs with two subsets. We had high overlapping among different runs and different subset for interactions and unique ceRNAs (Table 4).

Furthermore, we got the overlapping pairs for all subsets having 17,419 ceRNA interactions between unique 5223 ceRNAs. When we checked the distribution of overlapping ceRNAs in our inferred network, the degree distribution of these ceRNAs were much higher as compared to the overall degree distribution before filtering (p-value $< 2.10^{-16}$) suggesting that consistently inferred ceRNAs were the important ceRNAs highly involving in our inferred ceRNA network.

**Robustness of hyperparameter selection:** To show the hyperparameter selection in Crinet is robust to hyperparameter selection, we modified hyperparameters in ceRNA inference including network deconvolution step, and analyzed the results.

To show the robustness of ceRNA inference, we slightly changed hyperparameters eliminating more interactions with respect to one hyperparameter each time keeping similar number of interactions. The hyperparameters we used were hypergeometric p-value for common miRNA regulator among ceRNA pair, partial correlation and partial correlation bootstrapping threshold, and collective regulation correlation and its bootstrapping correlation, and we eliminated approximately one-third of the interactions concerning only one threshold for each run. Then we checked the overlapping interactions for them before network deconvolution filtering to see the effect. We got the overlapping of three different runs changing only one hyperparameter each time, and they had high overlapping having 17,419 interactions between unique 5223 ceRNAs, still keeping half of the interactions the same with highly-overlapping ceRNAs. (Table 6). This analysis suggests that even each hyperparameter had significant effect on the results, they consistently kept similar interactions with high number of unique overlapped ceRNAs.

To analyze that selection of ranking score in the incorporated network deconvolution algorithm does not change results much, we tried several values, and evaluated the results with LINCS data in a same way with pairwise ceRNA inference. Since the incorporated network deconvolution algorithm [2] gives a ranking score for interactions, but this ranking score is not showing exact significant, we had a wide range for ranking selection, but still the evaluation of obtained network was around the same accuracy (Table 5). These analysis showed that the network deconvolution step is important to eliminate indirect ceRNA interactions, but the selection of ranking did not significantly change Crinet results.

### A.4 Skipping individual steps made a substantial difference in inferred results

In this section, we assessed three major steps of ceRNA inference in our pipeline, namely: partial correlation filtering, collective regulation filtering, and exclusion of indirect ceRNA interactions. To examine the importance of each step, we skipped these steps to see how it would modify the inferred ceRNA network. Specifically, we kept all steps just skipping partial correlation filtering, referred to as “No CORR”. Since we had much more interaction at this step, we used -0.2 for collective regulation threshold, which is around 1.quartile of correlation distribution inferring 180,668 ceRNA interaction. As a second step, we just skipped collective regulation filtering, referred to as “No COLL”, and ran the pipeline similarly inferring 146,174 ceRNA interactions. Also, we got the results of our network before applying network deconvolution filtering, referred to as “No ND”. We also checked the skipped correlation and abundance filtering without network deconvolution. We called our results with whole steps as “Crinet”, and compare it in Table 1 with other runs using LINCS data that we used for network accuracy assessment.

Based on these results, skipping any step drastically changed the results. For the case “Crinet-no CORR”, the evaluation was based on a very limited number of perturbagens. On the other hand, even we had a slightly better result for “Crinet-no CORR no ND”, we had a huge number of inferred pairs, and a small number of perturbagen to evaluate with respect to the inferred number of pairs. For other cases, the accuracies of inferred networks were lower than the original results. Overall, these results suggest that the ceRNA interaction obtained from the modified network was not good at predicting gene expression change of its ceRNA partner, potentially inferring more false ceRNA interactions.

### B Supplementary Tables and Figures

**Table 1. Analysis to check the accuracy of inferred ceRNA interactions for modified pipeline using LINCS-L1000 shRNA-mediated gene knockdown experiment in breast cancer cell line** Table has interaction number, unique ceRNA number, and prediction accuracy of inferred network skipping different steps of Crinet. The values show the prediction accuracy of inferred network skipping different steps of Crinet along with the number of downstream ceRNAs whose ratios are smaller than 1 over all including the percentages. Crinet: Inferred results running whole pipeline; No ND: Skipping network deconvolution filtering; No CORR: Skipping expression correlation filtering; No COLL: Skipping collective regulation filtering.

| | Interaction# | ceRNA# | Overall Accuracy |
|---|---:|---:|---:|
| Crinet | 17,443 | 4,494 | 92/154=60% |
| No ND | 52,858 | 7,263 | 178/343=52% |
| No CORR | 59,620 | 7,541 | 12/21=57% |
| No CORR no ND | 180,668 | 10,895 | 44/71=62% |
| No COLL | 48,237 | 5,642 | 119/225=53% |
| No COLL no ND | 146,174 | 9,910 | 320/655=49% |

**Table 2. Analysis with miRNA transfection.** As negative control, we used non-inferred interactions using the same targets and miRNAs used in the corresponding step. Negative control shows the random preference being around 1 for each step when using non-inferred interactions with the same RNA and miRNAs in the corresponding step

| Step | Interaction Phase | Negative Control | Gene Phase | Negative Control |
|:---:|---:|---:|---:|---:|
| ALL | 6,248/4,932=1.3 | 5,964/5,980=1.0 | 90/74=1.2 | 69/95=0.7 |
| ALL40 | 613/403=1.5 | 11,118/9,862=1.1 | 97/59=1.6 | 74/82=0.9 |
| CORR | 183/112=1.6 | 7,778/7,155=1.1 | 71/37=1.9 | 48/60=0.8 |
| CORR+ABUN | 179/111=1.6 | 7,670/7,127=1.1 | 68/39=1.7 | 47/60=0.8 |

### References


1. E. P. Consortium et al. A user's guide to the encyclopedia of dna elements (encode). *PLoS biology*, 9(4), 2011.

2. S. Feizi, D. Marbach, M. Médard, and M. Kellis. Network deconvolution as a general method to distinguish direct dependencies in networks. *Nature biotechnology*, 31(8):726, 2013.

3. H. Han, J.-W. Cho, S. Lee, A. Yun, H. Kim, D. Bae, S. Yang, C. Y. Kim, M. Lee, E. Kim, et al. Trrust v2: an expanded reference database of human and mouse transcriptional regulatory interactions. *Nucleic acids research*, 46(D1):D380–D386, 2018.

4. C. Stark, B.-J. Breitkreutz, T. Reguly, L. Boucher, A. Breitkreutz, and M. Tyers. Biogrid: a general repository for interaction datasets. *Nucleic acids research*, 34(suppl_1):D535–D539, 2006.

**Table 3. Analysis with miRNA-protein expression anticorrelation.** Interaction phase shows the anticorrelated expressions of miRNA-protein target pairs with respect to not anticorrelated ones. Gene phase shows the anticorrelated expression of target with average of all miRNA regulators with respect to not-anticorrelated ones. (ratio is expected to be more than 1 to have anticorrelation preference. Higher is better). Negative control shows the random preference being around 1 for each step when using non-inferred interactions with the same RNA and miRNAs in the corresponding step. ALL: All obtained miRNA-target interactions; ALL40: Top 40% of ALL interactions; CORR: After expression correlation filtering on ALL40 interactions; CORR+ABUN: After abundance sufficiency filtering on CORR interactions.

| Step | Interaction Phase | Negative Control | Gene Phase | Negative Control |
|---:|---:|---:|---|---:|
| ALL | 38,505/33,243=1.2 | 52,690/53,849=1.0 | 95/106=0.9 | 106/95=1.1 |
| ALL40 | 2,720/2,688=1.0 | 80,157/76,993=1.0 | 107/91=1.2 | 93/105=0.9 |
| CORR | 751/370=2.0 | 36,855/34,459=1.1 | 115/50=2.3 | 74/91=0.8 |
| CORR+ABUN | 719/352=2.0 | 35,004/32,672=1.1 | 114/47=2.4 | 72/89=0.8 |

**Table 4. Reproducibility of Crinet with different subsets.** Overlapping interaction number and overlapping unique ceRNA number (in parentheses) are shown in the lower triangular; while overlapping percentages of the cases in row and in column with respect to the smaller number are shown in upper triangular.

| Case | Case-inferred | 1.Run 1.Set | 1.Run 2.Set | 2.Run 1.Set | 2.Run 2.Set |
|---|---:|---:|---:|---:|---:|
| 1.Run 1.Set | 8,156 (2,990) | - | 47% (82%) | 54% (81%) | 54% (86%) |
| 1.Run 2.Set | 18,093 (4,273) | 3,834 (2,445) | - | 58% (88%) | 66% (87%) |
| 2.Run 1.Set | 11,022 (3,559) | 4,399 (2,431) | 6,353 (3,122) | - | 51% (84%) |
| 2.Run 2.Set | 18,097 (4,210) | 11,878 (3,674) | 11,020 (4,441) | 5,576 (2,980) | - |

**Table 5. Analysis to check the accuracy of pipeline with different ranking of network deconvolution based on LINCS-L1000 (MCF7) dataset.**

| | Interaction# | 96h Timepoint | 144h Timepoint | Overall |
|:-:|---:|---:|---:|---:|
| Top 10% | 5,286 | 22/31=71% | 13/30=43% | 35/61=57% |
| Top 15% | 7,929 | 30/42=71% | 24/43=56% | 54/85=64% |
| Top 20% | 10,572 | 35/50=70% | 27/50=54% | 62/100=62% |
| Top 25% | 13,214 | 40/60=67% | 31/61=51% | 71/121=59% |
| Top 15,000 | 15,000 | 43/68=63% | 35/68=51% | 78/136=57% |
| Top 30% | 15,857 | 45/72=62% | 39/72=54% | 84/144=58% |
| **Top one third** | **17,443** | **48/77=62%** | **44/77=57%** | **92/154=60%** |
| Top 35% | 18,500 | 49/79=62% | 44/78=56% | 93/157=59% |
| Top 20,000 | 20,000 | 54/86=63% | 46/84=55% | 100/170=59% |
| Ranking>0.7 | 20,423 | 55/88=62% | 47/86=55% | 102/174=59% |
| Top 40% | 21,143 | 55/93=59% | 47/91=52% | 102/184=55% |
| Top 25,000 | 25,000 | 59/100=59% | 48/98=49% | 107/198=54% |
| Ranking>3rd quartile | 26,428 | 61/104=59% | 49/103=48% | 110/207=53% |
| Top 50% | 26,429 | 61/104=59 | 49/103=48% | 110/207=53% |
| Top 60% | 31,715 | 70/125=56 | 55/121=45% | 125/246=51% |
| Top 70% | 37,001 | 78/136=57 | 72/132=55% | 150/268=56% |
| Top 80% | 42,284 | 84/146=58 | 80/144=56% | 164/290=57% |

**Table 6. Analysis to show the robustness to different hyperparameter selection of Crinet.** The hyperparameters are hypergeometric p-value for common miRNA regulator among ceRNA pair (referred to as thr1), partial correlation and partial correlation bootstrapping threshold (referred to as thr2), and collective regulation correlation and its bootstrapping correlation (referred to as thr3), and we eliminated approximately one-third of the interactions with respect to only changing one threshold for each run. The values shows the overlapping of three different runs changing only one hyperparameter

| | thr1 | thr2 | thr3 | Interaction # | Overlap | ceRNA # | Overlap |
|---|---|---|---|---|---|---|---|
| Run1 | **0.0011** | 0.55 | -0.01 | 35,509 | | 6,636 | |
| Run2 | 0.01 | **0.58** | -0.01 | 37,107 | 17,419 | 6,175 | 5,223 |
| Run3 | 0.01 | 0.55 | **-0.03** | 36,729 | | 6,204 | |

**Table 7. GO Slim terms from the biological process ontology enriched for the inferred ceRNA groups with coding genes. We put the GO Slim terms with at least half of the genes in the group. Group indicates the group we analyzed deeply shown in Figure 4). Definition is the name of GO term. Gene ratio is the fraction of inferred genes in the term to total applicable genes in the term. We used the tool named GO Term Mapper from URL** `https://go.princeton.edu/cgi-bin/GOTermMapper`

| Group # | GO term ID | Definition | Gene Ratio |
|---|---|---|---|
| 1 | GO:0007049 | cell cycle | 6/6 |
| 1 | GO:0006950 | response to stress | 4/6 |
| 1 | GO:0006259 | DNA metabolic process | 4/6 |
| 1 | GO:0051276 | chromosome organization | 4/6 |
| 1 | GO:0034641 | cellular nitrogen compound metabolic process | 4/6 |
| 1 | GO:0007059 | chromosome segregation | 3/6 |
| 1 | GO:0009058 | biosynthetic process | 3/6 |
| 1 | GO:0051301 | cell division | 3/6 |
| 1 | GO:0007010 | cytoskeleton organization | 3/6 |
| 2 | GO:0002376 | immune system process | 1/2 |
| 2 | GO:0030154 | cell differentiation | 1/2 |
| 3 | GO:0002376 | immune system process | 3/4 |
| 3 | GO:0048870 | cell motility | 3/4 |
| 3 | GO:0007165 | signal transduction | 3/4 |
| 3 | GO:0040011 | locomotion | 3/4 |
| 3 | GO:0008219 | cell death | 2/4 |
| 3 | GO:0007155 | cell adhesion | 2/4 |
| 3 | GO:0048856 | anatomical structure development | 2/4 |
| 4 | GO:0002376 | immune system process | 3/4 |
| 4 | GO:0006950 | response to stress | 2/4 |

**Table 8. GO Slim terms from the biological process ontology enriched for the inferred ceRNAs with applicable genes. We put the GO Slim terms with at least 100 genes in the term. Definition is the name of GO term. Gene ratio is the fraction of inferred genes in the term to total applicable genes in the term. We used the tool named GO Term Mapper from URL** `https://go.princeton.edu/cgi-bin/GOTermMapper`

| GO term ID | Definition | Gene Ratio |
|---|---:|---:|
| GO:0034641 | cellular nitrogen compound metabolic process | 1076/2888 |
| GO:0009058 | biosynthetic process | 1008/2888 |
| GO:0007165 | signal transduction | 941/2888 |
| GO:0048856 | anatomical structure development | 929/2888 |
| GO:0006810 | transport | 723/2888 |
| GO:0006950 | response to stress | 701/2888 |
| GO:0006464 | cellular protein modification process | 674/2888 |
| GO:0030154 | cell differentiation | 646/2888 |
| GO:0002376 | immune system process | 631/2888 |
| GO:0022607 | cellular component assembly | 448/2888 |
| GO:0008219 | cell death | 375/2888 |
| GO:0009056 | catabolic process | 369/2888 |
| GO:0007049 | cell cycle | 353/2888 |
| GO:0040011 | locomotion | 341/2888 |
| GO:0007155 | cell adhesion | 331/2888 |
| GO:0016192 | vesicle-mediated transport | 331/2888 |
| GO:0048870 | cell motility | 311/2888 |
| GO:0065003 | protein-containing complex assembly | 287/2888 |
| GO:0042592 | homeostatic process | 244/2888 |
| GO:0007267 | cell-cell signaling | 230/2888 |
| GO:0044281 | small molecule metabolic process | 229/2888 |
| GO:0048646 | anatomical structure formation involved in morphogenesis | 220/2888 |
| GO:0000278 | mitotic cell cycle | 214/2888 |
| GO:0051276 | chromosome organization | 214/2888 |
| GO:0007010 | cytoskeleton organization | 211/2888 |
| GO:0000003 | reproduction | 200/2888 |
| GO:0009790 | embryo development | 187/2888 |
| GO:0006259 | DNA metabolic process | 185/2888 |
| GO:0055085 | transmembrane transport | 169/2888 |
| GO:0006629 | lipid metabolic process | 167/2888 |
| GO:0044403 | symbiont process | 156/2888 |
| GO:0000902 | cell morphogenesis | 152/2888 |
| GO:0040007 | growth | 151/2888 |
| GO:0051301 | cell division | 127/2888 |
| GO:0061024 | membrane organization | 124/2888 |
| GO:0050877 | nervous system process | 119/2888 |
| GO:0030198 | extracellular matrix organization | 106/2888 |

**Table 9. GO Slim terms from the biological process ontology enriched for the high-degreed ceRNAs ($> 50$ degrees) with applicable genes. We put the GO Slim terms with at least 10% of the genes in the term. Definition is the name of GO term. Gene ratio is the fraction of inferred genes in the term to total applicable genes in the term. We used the tool named GO Term Mapper from URL** `https://go.princeton.edu/cgi-bin/GOTermMapper`

| GO term ID | Definition | Gene Ratio |
|---|---:|---:|
| GO:0007165 | signal transduction | 26/45 |
| GO:0034641 | cellular nitrogen compound metabolic process | 17/45 |
| GO:0002376 | immune system process | 17/45 |
| GO:0006810 | transport | 15/45 |
| GO:0006464 | cellular protein modification process | 15/45 |
| GO:0022607 | cellular component assembly | 14/45 |
| GO:0009058 | biosynthetic process | 13/45 |
| GO:0008219 | cell death | 13/45 |
| GO:0048856 | anatomical structure development | 13/45 |
| GO:0006950 | response to stress | 12/45 |
| GO:0065003 | protein-containing complex assembly | 11/45 |
| GO:0030154 | cell differentiation | 10/45 |
| GO:0009056 | catabolic process | 10/45 |
| GO:0008283 | cell proliferation | 9/45 |
| GO:0048870 | cell motility | 9/45 |
| GO:0040011 | locomotion | 9/45 |
| GO:0042592 | homeostatic process | 8/45 |
| GO:0007155 | cell adhesion | 8/45 |
| GO:0016192 | vesicle-mediated transport | 7/45 |
| GO:0044403 | symbiont process | 5/45 |
| GO:0061024 | membrane organization | 5/45 |
| GO:0007049 | cell cycle | 5/45 |

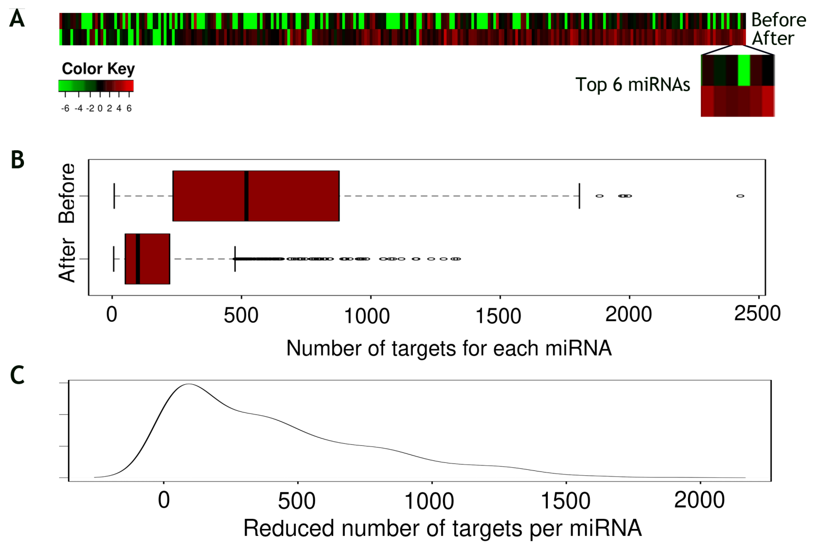


**Figure 1. Plot showing the analysis of abundance sufficiency filtering for miRNA-target interactions.** A. Heatmap showing median log expression of top 25% miRNAs with the highest and lowest number of targets in miRNA-target interaction set obtained before and after abundance filtering. The miRNAs are ranked from right to left based on target numbers where the rightmost one is the miRNA with the highest target number. B. Boxplot showing the number of targets for each miRNA in the miRNA-target interaction set obtained before and after abundance filtering. C. Density plot showing the reduced number of targets for each miRNA in the miRNA-target interactions when miRNAs are filtered based on abundance sufficiency.

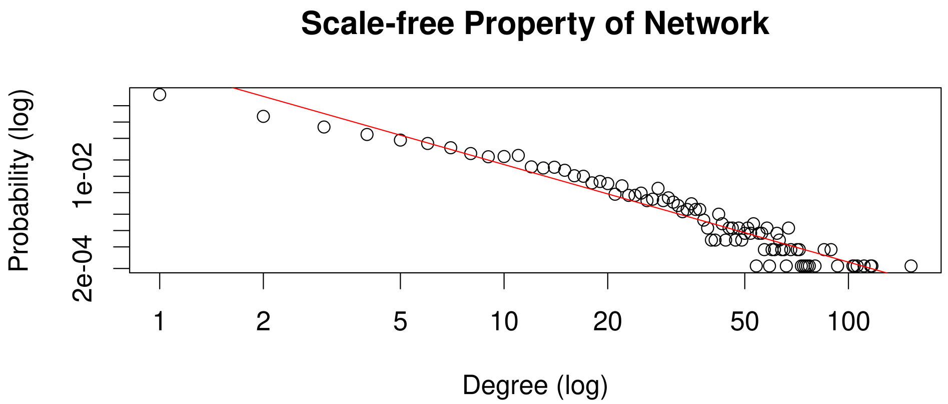


**Figure 2. Scale-free property of the inferred network.** Computing inferred network degree probability distribution function, and following the power-law rule, we fitted linear regression for the log of ceRNA's degree probability to the log of ceRNA's degree. The plot for the inferred ceRNA network had a negative slope with high fitness ($R^2 = 0.93$), indicating that the inferred ceRNA network was scale-free.

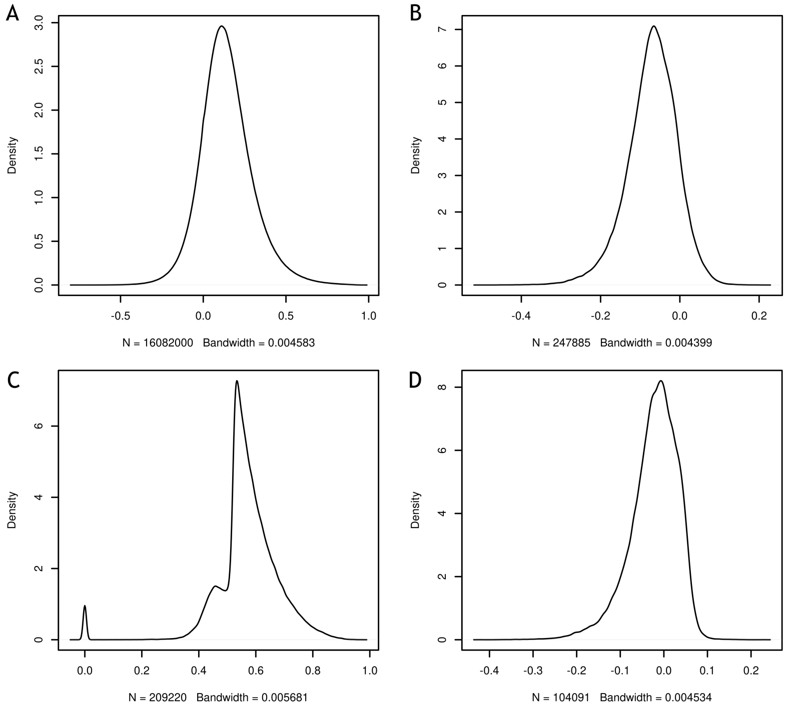


**Figure 3. Density plots showing the correlation distribution for filtering steps** A. Density plot showing the partial expression correlation distribution excluding copy number aberration effect for candidate ceRNA pairs having at least one common regulator with significant overlap (partial correlation filtering). B. Density plot showing the correlation distribution among the sum of Effective Regulation scores and the sum of the expression of each pair (collective regulation filtering) following partial correlation filtering. C. Density plot showing the maximum threshold for partial expression correlation excluding copy number aberration to keep the ceRNA pair satisfying 99 out of 100 calculations with replacement (partial correlation bootstrapping) following collective regulation filtering. D. Density plot showing the maximum threshold for correlation among the sum of Effective Regulation scores and the sum of the expression to keep the ceRNA pair satisfying 99 out of 100 calculations with replacement following partial correlation bootstrapping.

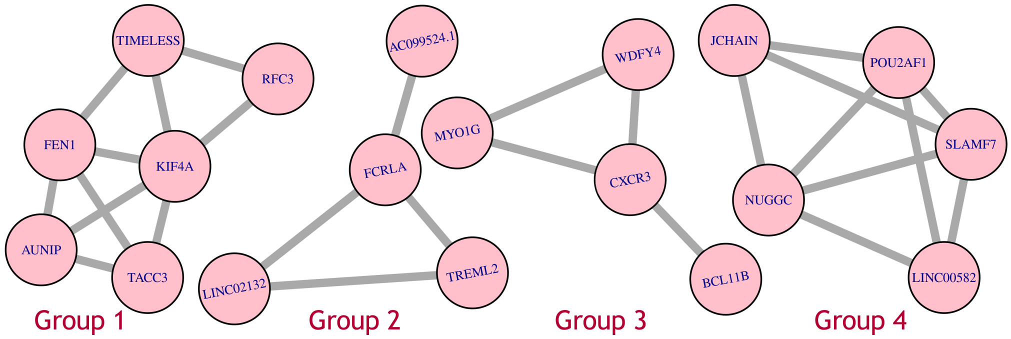


**Figure 4. Deeply analyzed inferred ceRNA groups**
